# Glutamatergic Neurons in caudal Pontine Reticular Nucleus (PRNc) Mediate Backward Locomotion in mice

**DOI:** 10.64898/2026.10.22.761180

**Authors:** Michelle Boyoon Seo, Junxiang J. Huang, Guang-Wei Zhang, Kyle Voong, Huizhong Whit Tao, Li I. Zhang

## Abstract

Locomotion is essential for animals to adaptively navigate their environment. However, despite extensive investigation into neural mechanisms underlying forward movement, the brain circuits controlling backward locomotion, a critical motor behavior for navigating confined spaces and avoiding threats, remain poorly understood. Here, using a specially designed behavioral paradigm for backward locomotion in mice, we find that glutamatergic neurons in the caudal pontine reticular nucleus (PRNc) are selectively activated during backward, but not forward, locomotion. Activity increase of these neurons is temporally aligned with backward movement, with its amplitude correlated with speed. Optogenetic manipulations further demonstrate that these neurons are both sufficient and necessary for initiating and sustaining backward walking. Moreover, anatomical tracing reveals that the PRNc broadly integrates bilateral cortical and subcortical inputs and projects extensively to brain-wide and spinal cord networks. These findings suggest PRNc as a previously unrecognized brainstem center in direction-specific locomotor control circuits. Key findings of this work and our initial identification of this brainstem locomotor node were presented at the Society for Neuroscience Annual Meeting in November, 15^th^, 2025 (SfN Neuroscience 2025, San Diego, CA; Program No. PSTR027.02)^1^.

## Introduction

Locomotion is critical for an animal’s survival, enabling navigation, foraging, exploration, and threat avoidance. To flexibly adapt to varying environmental demands and internal motivations, animals must integrate sensory inputs, make context-dependent decisions and execute the most appropriate locomotor strategy such as modulating speed, stopping, turning, and moving forward or backward ^2–5^. Although previous studies have predominantly focused on forward locomotion ^2–7^, backward locomotion is an equally important behavior observed across species. For example, burrowing animals navigate tight tunnels by backing out, predators reposition during pursuit, and humans use backward movements in sports and tactical settings. Despite its functional relevance, backward locomotion has received comparatively little attention especially in mammals due to its context-dependent nature. Consequently, brain circuits specifically underlying backward movement remain poorly understood, despite some studies in the spinal cord ^3–5,8,9^. Given the ethological significance and ubiquity of backward movements, identifying dedicated neural circuits governing this behavior is critical to fully elucidate the functional organization of motor control systems.

Recent work in non-mammalian model organisms has begun to reveal dedicated neural substrates for backward locomotion. In the fruit fly, specialized neuronal populations, such as Moonwalker Descending Neurons (MDNs) and Moonwalker Subesophageal Zone neurons (MooSEZs) have been shown to selectively mediate backward walking ^10,11^. Similarly, in lower vertebrates like the lamprey, certain reticulospinal neurons exhibit direction-specific activity patterns, with subsets exclusively active during backward swimming ^12^. These studies suggest the existence of dedicated neural modules for backward locomotion across diverse species.

Additionally, evidence across species suggests that forward and backward locomotion may involve partially segregated spinal and supraspinal mechanisms rather than a single bidirectional circuit ^8,9,12^. However, in mammalian brain, it remains unclear whether any specific circuits or molecularly defined neuronal populations specifically control backward locomotion. Recent studies have shown that the mammalian brainstem contains modular neuronal populations specialized for controlling various forms of motor behaviors including forward running, locomotor arrest, turning, and forelimb control ^4,3,13–19^. These findings raise the possibility that the mammalian brainstem might also harbor dedicated circuits specifically governing backward locomotion.

To address this question, we developed a behavioral paradigm in mice to reliably induce backward locomotion, which allows us to systematically investigate the locomotor control at behavioral and neural circuit levels. Combining with activity mapping, calcium imaging, optogenetic manipulations, and anatomical tracing, we identified glutamatergic neurons in the caudal pontine reticular nucleus (PRNc) as a direction-selective neural population that is specifically activated during backward but not forward movement. Functional analyses confirmed that these neurons are sufficient and necessary to initiate and sustain backward locomotion, while anatomical tracing revealed their integration within broader brainstem, cortical, and spinal networks. Collectively, our findings establish the PRNc as a dedicated brainstem center governing backward locomotion and provide new insight into the modular organization of direction-specific locomotor control circuits. Furthermore, these results may offer new perspectives on clinical conditions characterized by impaired motor coordination, gait instability, or directional movement deficits, such as Parkinson’s disease, stroke, or spinal cord injury, which may help with developing targeted therapeutic strategies for motor rehabilitation and recovery.

## Results

### Kinematics of backward locomotion

To establish a behavioral foundation for our neural analyses of backward movement, we first compared the kinematics of backward walking (BW) in three scenarios― spontaneous (Spont. BW), trained (Trained BW), and defensive (Def. BW) backward walking― relative to forward walking (FW) (**Figure 1A-I**). We designed a transparent, straight and narrow corridor for mouse locomotion that prevents turning, and recorded a bottom-view video. We then applied DeepLabCut ^20,21^, a deep learning-based pose estimation toolbox, to track the nose, neck, tail-base, and all four paws of the mouse—enabling precise measurements of velocity and gait metrics (see Methods). We began by characterizing the spontaneous backward locomotion in the corridor (**Figure 1A**). Mice would occasionally enter the corridor and, unable to turn, back out in a series of short, segmented bouts, as shown by the plot of the tail-base movement velocity (**Figure 1B**). It has been reported that a sudden visual threat could trigger a backward retreat ^22–24^. To mimic this, we presented a novel object suddenly dropping in front of—but not touching— the mouse. This triggered a defensive response associated with brief, high-speed backward locomotion across all animals (**Figure 1E-F**). However, with repeated exposures, mice typically habituated to the stimulus and the response became less frequent. Because both spontaneous and defensive backward movements were rare and relatively brief (**Figure 1J**), we devised and validated a novel trained task that elicited repeated long bout backward walking in mice actively seeking water reward (**Figure 1G-H**), enabling a large number of trials in each animal for robust movement analysis (**Figure 1I-M**).

**Figure 1.**
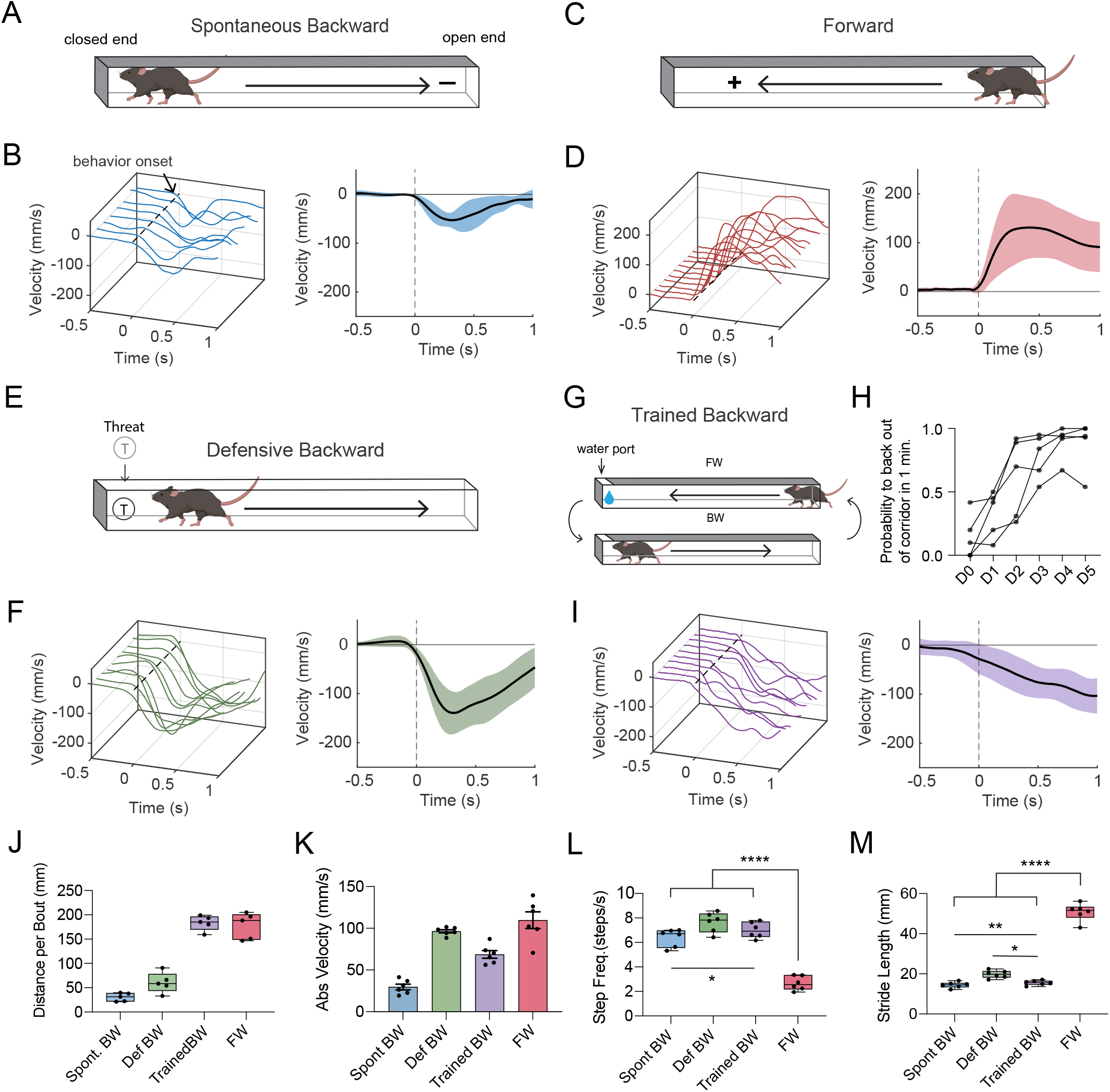
Characterization of backward movement in different contexts. (A) Schematic spontaneous backward locomotion in a corridor assay. Arrow marks the direction of locomotion. (B) Left, example velocity traces aligned to the onset of spontaneous backward locomotion (marked by vertical dotted line). Velocity was calculated based on movements of the tail base. Negative values denote backward direction. Right, population average of velocity across all bouts from five animals. Shade denotes ±1 SD. (C-D) Similar to (A-B) but for forward locomotion. n = 5 mice. (E-F) Similar to (A-B) but for defensive backward locomotion. n = 5 mice. (G) Schematic training of a mouse to enter a corridor, receive water reward and back out. (H) Probability of backing out of the corridor within 1 min across training days. Data points for the same animals are connected. (I) Plots of example velocity traces and the population average. n = 5 mice. (J) Comparison of mean distance per bout in different contexts. (K) Comparison of mean velocity (absolute value) within a 1-second window following movement onset across four conditions (n = 5 mice per condition): spontaneous backward (Spont. BW), trained backward (Trained BW), defensive backward (Def. BW), and forward (FW) locomotion. *p < 0.05, **p < 0.01, ***p < 0.001, ****p < 0.0001, ns, not significant, p > 0.05, one-way ANOVA with Tukey’s post hoc test. Bar = SD. (L) Comparison of mean step frequency across conditions. Only statistically different pairs are annotated. *p < 0.05, ****p < 0.0001 for Spont. BW vs FW, Trained BW vs. FW, Def. BW vs. FW, one-way ANOVA with Tukey’s post hoc test. (M) Comparison of mean stride length across all conditions. *p < 0.05, **p < 0.01, ****p < 0.0001 for Spont. BW vs FW, Trained BW vs. FW, Def. BW vs. FW, one-way ANOVA with Tukey’s post hoc test.

Analysis of movements of the tail-base revealed that spontaneous BW and defensive BW were much briefer than the trained BW and FW, as shown by the differences in the mean distance per bout (**Figure 1J**). Additionally, locomotion speeds of both the spontaneous and trained BW were significantly slower than FW, whereas the velocity of defensive BW was similar to FW (**Figure 1K**). These results demonstrate the adaptive nature of backward locomotion, showing its ability to overcome biomechanical constraints and achieve speeds comparable to forward locomotion.

In addition to the overall velocity, backward walking displayed a distinct limb cycle compared to forward movement. Step frequency, measured only during active stepping, was significantly higher in all BW contexts than forward walking (**Figure 1L**). Among the different BW contexts, defensive BW exhibited the highest step frequency (**Figure 1L**), which is consistent with its rapid acceleration to escape. Conversely, stride length was markedly shorter in all BW contexts compared to FW (**Figure 1M**), consistent with previous reports ^9,25–27^. Spontaneous and trained BW bouts showed similar stride lengths, whereas defensive BW displayed slightly longer strides than the other BW contexts (**Figure 1M**).

Together, our behavioral analyses demonstrate that mice can flexibly adjust their BW speed by modulating both step frequency and stride length in tandem, to suit varying demands. In low arousal contexts, a modest increase in step frequency combined with shorter strides suffices for slow exploratory reversals, whereas under defensive conditions a higher step frequency along with longer strides enables rapid retreats with speeds comparable to forward movement.

### PRNc neurons activated by repeated backward walking

To identify which brain regions are recruited by backward locomotion, we employed TRAP2::Ai14 reporter mice, allowing permanent labeling of neurons active within a time window following 4-hydroxytamoxifen (4-OHT) administration ^28,29^. Water restricted mice were trained for five days to seek water reward by navigating through a narrow corridor (**Figure 2A**)., They learned to perform forward-only (with both corridor ends open) or forward-followed-by-backward (with one corridor end closed) navigation depending on the context (see Methods). On the test day, the trained animals were randomly assigned into a FW or BW group. Following 4-OHT injection, animals performed 50 trials of either forward walking only (FW group) or alternating forward and backward episodes (BW group) (**Figure 2A**). To analyze TRAPed (i.e. tdTomato-labeled) neurons, we focused on brainstem nuclei known to govern locomotor control such as the gigantocellular reticular nucleus (GRN), magnocellular reticular nucleus (MARN) and intermediate reticular nucleus (IRN) ^3,17,30^, as well as midbrain centers implicated in defensive responses including the periaqueductal gray (PAG), superior colliculus (SC), and parabrachial nucleus (PBN) ^31–34^. We also examined regions showing dense labeling in both FW and BW conditions such as the tegmental reticular nucleus (TRN), pontine gray (PG), and pontine central gray (PCG). Analysis of these areas revealed that there was an _∼_3-fold increase in the number of tdTomato neurons in PRNc in the BW relative to FW condition (**Figure 2B-C**). To further characterize the cell type of the TRAPed neurons in the PRNc, RNAscope staining was performed using probes for vesicular glutamate transporter 2 (Vglut2), which is a marker for excitatory glutamatergic neurons. We found that neurons activated by repeated BW bouts were primarily excitatory, as 76.2 ± 4.1% of the TRAPed neurons were Vglut2+ (**Figure 2D-E**). In sum, our results reveal that glutamatergic neurons in the PRNc are preferentially recruited by backward walking, laying the groundwork for targeted calcium imaging and manipulations to examine a causal role of these neurons in backward locomotion control.

**Figure 2.**
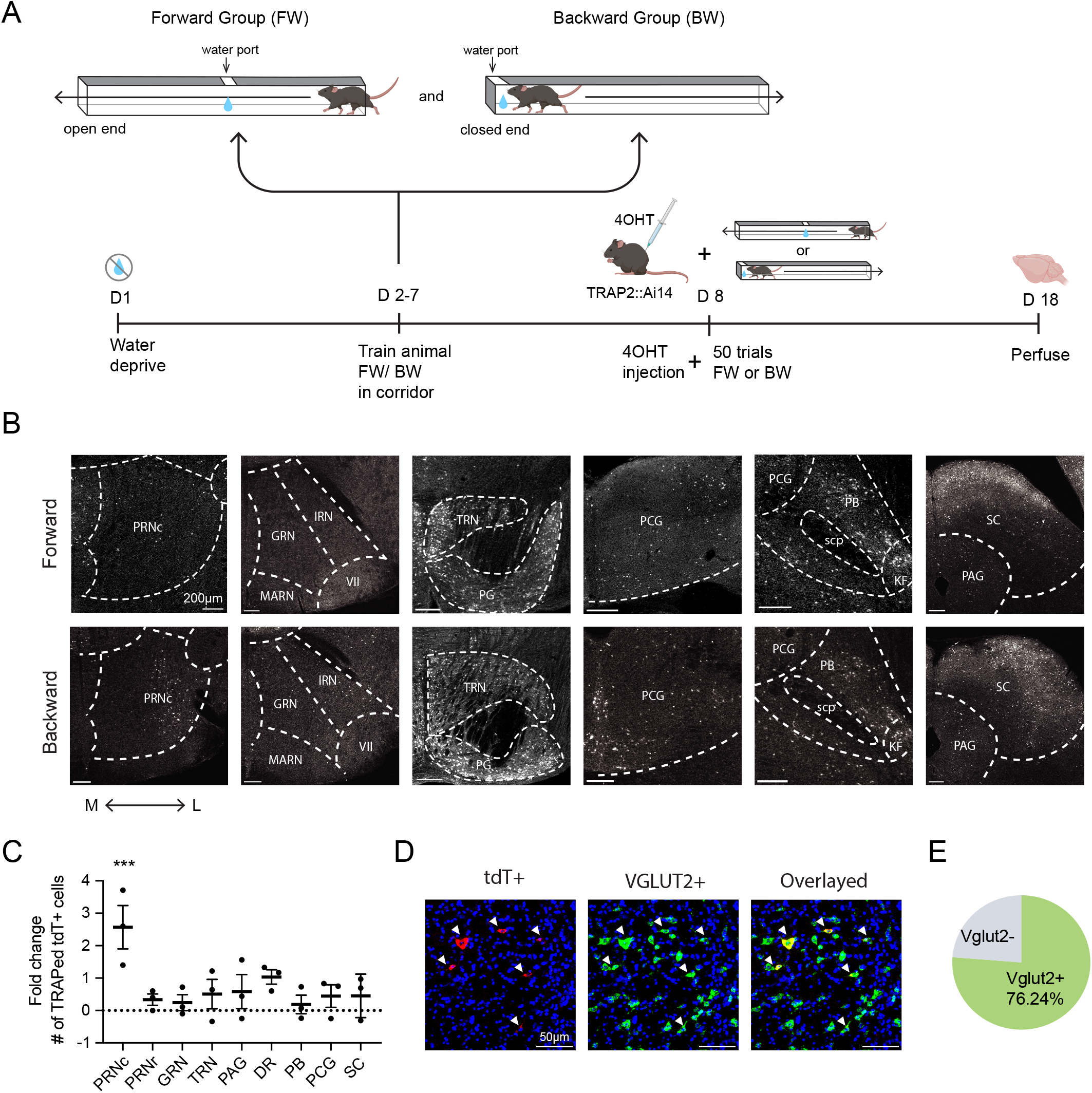
Screening of brainstem regions activated after repeated backward movements. (A) Schematic of timeline for c-Fos labeling. (B) Representative confocal images of tdTomato+ (i.e. c-Fos expressing) neurons in coronal sections from FW (top row) and BW (bottom row) groups. Dashed outlines demarcate anatomical nuclei: caudal pontine reticular nucleus (PRNc); intermediate reticular nucleus (IRN); gigantocellular reticular nucleus (GRN); facial nucleus (VII) and magnocellular reticular nucleus (MARN); tegmental reticular nucleus (TRN) and pontine gray (PG); pontine central gray (PCG); parabrachial nucleus (PB), superior cerebellar peduncle (scp) and Koelliker-Fuse nucleus (KF); superior colliculus (SC) and periaqueductal gray (PAG). Medial (M) → Lateral (L) orientation is indicated at the bottom. Scale bar = 200 µm. (C) Fold change in the number of tdTomato+ (tdT) TRAPed cells in each nucleus for the BW cohort relative to the FW cohort. Data are plotted as mean ± SEM with individual dots representing one animal (n = 3 mice per group). A significant increase in TRAPed neurons was observed in PRNc (***p = 0.0008, one sample t test vs. 0). (D) Representative RNAscope images in the PRNc showing tdT expression (red, left), Vglut2 expression (green, middle), and overlaid (right). Arrowheads mark co-labeled cells (yellow). Scale bar = 50 µm. (E) Proportion of TRAPed neurons in PRNc that co-express Vglut2 (green) versus those that do not (gray).

### Ca^2+^ dynamics of PRNc glutamatergic neurons during backward movement

To further investigate the recruitment of PRNc glutamatergic neurons by backward locomotion, we virally expressed the Cre-dependent Ca^2+^ indicator, GCaMP7s, in PRNc glutamatergic neurons using Vglut2-Cre mice (**Figure 3A**). With fiber photometry, bulk fluorescence signals were recorded as animals moved in and out of the corridor (see Methods). The recorded Ca^2+^ events were subsequently aligned to the onsets of BW or FW bouts, identified based on the mouse’s velocity (**Figure 3B**). In the majority of trials, Ca^2+^ activity increased and peaked following the onset of BW (**Figure 3C-E**), whereas FW was associated with a slight decrease in fluorescence intensity (**Figure 3F-H**), with both responses differing from the baseline (0-2 s window area under curve: BW, 1.78 vs. 0, p < 0.0001; FW,-0.39 vs. 0, p = 0.385, one-sample t-test). Comparison of the average peak ΔF/F values from all animals revealed a significant difference between BW and FW (**Figure 3I**). Furthermore, correlation analysis revealed a negative relationship between levels of Ca^2+^ activity and locomotion velocity (**Figure 3J**). This suggests that PRNc glutamatergic neuron activity is dynamically modulated by the direction and velocity of locomotion, with higher activity of these neurons associated with faster backward movements.

**Figure 3.**
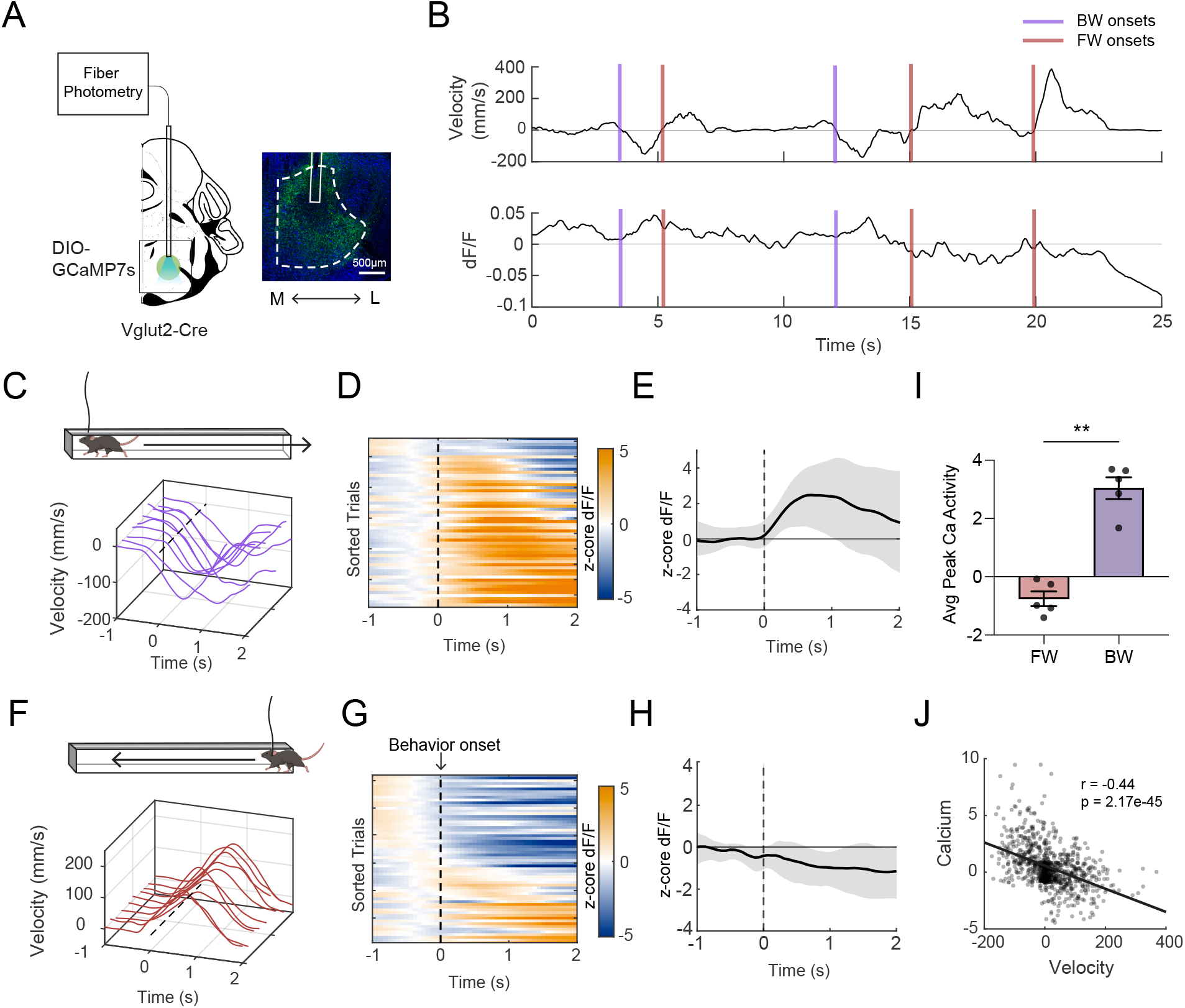
Ca^2+^ dynamics of PRNc glutamatergic neurons during forward and backward movements. (A) Left, schematic for viral injection and fiber photometry recording. Right, example image for GCaMP expression at the injection site (green: GCaMP; blue: Nissl). Dashed outline marks the PRNc. Scale bar = 500 µm. (B) Example traces of locomotor velocity (top) and fluorescence change (ΔF/F, bottom) in the same time window. Colored vertical lines mark the onsets of BW (purple) and FW (red) bouts. (C) Representative velocity traces from ten BW trials, aligned to the onset of behavioral bouts (time 0, dashed line). (D) Heatmap of z scored ΔF/F responses in PRNc Vglut2 neurons for all backward trials (n = 55 trials from 5 mice), sorted by the latency to peak velocity (color bar = z score). (E) Population average of z scored ΔF/F (black line) ± 1 SD (shaded), aligned to movement onset. (F-H) Similar to (C-E) but for FW trials (n = 61 trials from 5 mice). (I) Average peak z scored ΔF/F amplitude in the 0–2 s window following locomotion onset for FW versus BW trials (n = 5 mice). **p = 0.0015, paired t test. Bar represents SEM. (J) Plot of mean ΔF/F versus mean velocity computed in individual moving windows (window size = 5 frames) over the 0–2 s post-onset interval. Each point represents one window, and the solid line shows the linear regression (*r* = –0.44, p = 2.17 × 10).

### Optogenetic activation of PRNc glutamatergic neurons drives backward walking

To test whether PRNc glutamatergic neurons can drive backward locomotion, we expressed Cre-dependent channelrhodopsin-2 (ChR2) in Vglut2-Cre mice and delivered trains of 470 nm LED stimulation at varying frequencies (5, 10, or 20 Hz, 1-sec duration, 25-ms pulses) in the corridor assay (**Figure 4A**). The optical stimulation reliably triggered backward walking in a frequency-dependent manner, producing mean velocities of −52.40 ± 10.92 mm/s, −115.80 ± 19.06 mm/s, and −190.20 ± 24.51 mm/s at 5, 10, and 20 Hz, respectively (**Figure 4B–E**). These results demonstrate that activation of PRNc glutamatergic neurons is sufficient to drive backward locomotion and that increasing the activation frequency proportionally enhances the speed of backward movement. Gait analysis further revealed that the increase in BW speed was primarily due to an elevated step frequency (**Figure 4F**), while stride length did not significantly differ among different stimulation frequencies (**Figure 4G**). However, the variability of stride length was reduced at the highest testing frequency (5 Hz vs 20 Hz, variances 38.28 mm² vs 28.70 mm², * p = 0.041; 5 Hz vs 10 Hz, variances 38.28 mm² vs 35.04 mm², p = 0.508; 10 Hz vs 20 Hz, variances 35.04 mm² vs 28.70 mm², p = 0.132, pairwise F-test of variance) (**Figure 4H**).

**Figure 4.**
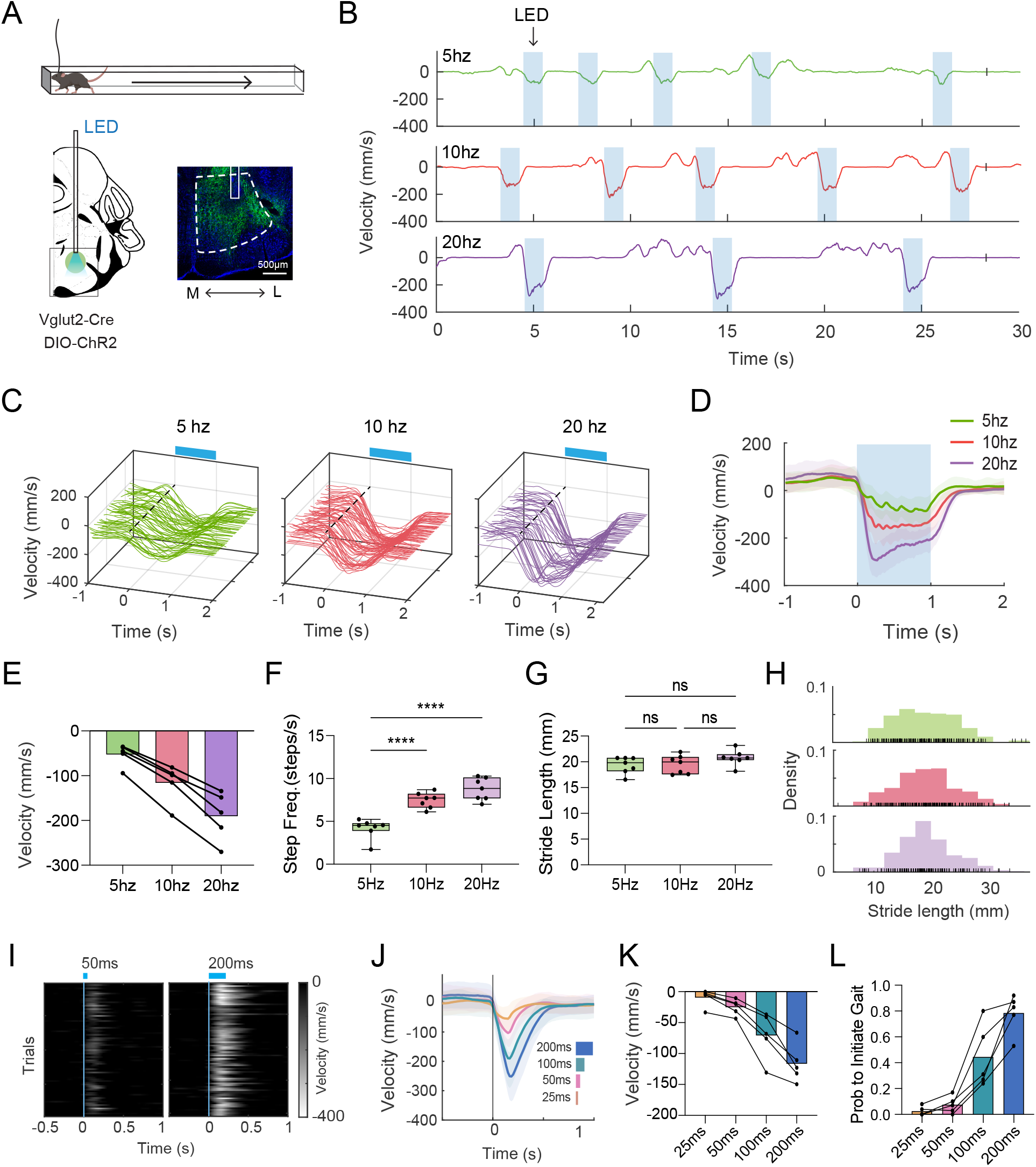
Activation of PRNc glutamatergic neurons drives backward locomotion. (A) Left, schematic of viral injection and optogenetic activation. Right, example image for ChR2-eYFP expression (green) and the fiber tract (white box) in PRNc. Scale bar = 500 µm. (B) Representative velocity traces during 1-s LED stimulation (blue shade) at 5 Hz (top, green), 10 Hz (middle, red), and 20 Hz (bottom, purple). (C) Individual velocity traces for all stimulation trials, aligned to the LED onset (time 0, dashed line), at different stimulation frequencies (same color code). Blue bar depicts the stimulation duration. (D) Group averaged velocity profiles at different stimulation frequencies (mean ± SD). n = 62, 78, 49 trials from 5 mice. (E) Mean velocity during the 1-s stimulation at different stimulation frequencies. Each dot represents one animal (n = 5). Data points for the same animals are connected. Column represents average across animals. *p = 0.042, ***p = 0.0007, one-way ANOVA with Tukey’s post hoc test. (F) Raster plot of stance onsets of right hindlimb during the 1-s LED stimulation (blue line) at different stimulation frequencies. Each tick mark denotes the moment the paw made full ground contact, aligned to the first stance event. (G) Mean step frequency (steps/s) averaged across all four limbs at different stimulation frequencies (n = 5 mice). Box spans the interquartile range with a central median line; whiskers indicate the minimum and maximum values. ****p < 0.0001, one-way ANOVA with Tukey’s post hoc test. (H) Mean stride length (mm) averaged across all limbs at different stimulation frequencies (n = 5 mice). Conventions as in (G); ns, not significant, p > 0.05, one-way ANOVA with Tukey’s post hoc test. (I) Stride length distributions at different stimulation frequencies. Histograms show density across animals, with each individual stride marked as a tick along the x axis. (J) Individual velocity traces for 50-ms and 200-ms single pulse stimulation, aligned to LED onset (dashed line). Blue bar indicates the duration of light pulses. (K) Group averaged velocity (mean ± SD) at different pulse durations: 25 ms (orange), 50 ms (pink), 100 ms (green), and 200 ms (blue). (L) Mean velocity during the 0–0.5 s post onset window for each pulse duration (n = 5 mice). *p < 0.05, **p < 0.01, one-way ANOVA with Tukey’s post hoc test. (M) Probability of evoking a backward walk (≥2 backward steps) after single pulse stimulation. *p < 0.05, ***p < 0.001, one-way ANOVA with Tukey’s post hoc test.

We further tested single pulse LED stimulation of varying durations (25, 50, 100, and 200 ms) (**Figure 4I-L**). Shorter pulses typically induced subtle backward shift of the body rather than complete stepping, suggesting an anticipatory postural adjustment prior to full gait initiation **(Figure 4I-K)**. Increasing the stimulus duration gradually enhanced the backward movement velocity as well as the probability of backward gait, as defined by at least one complete cycle of backward stepping for both hind limbs (**Figure 4K-L**). Together, these results suggest that PRNc glutamatergic neurons can flexibly control motor output, initiating either a postural adjustment or full backward locomotion depending on the duration and frequency of activation.

### Optogenetic inhibition of PRNc glutamatergic neurons impairs backward movement

To determine whether PRNc glutamatergic neurons are required for backward locomotion, we expressed the inhibitory opsin, GtACR, in these neurons using Vglut2-Cre mice and applied continuous LED light illumination during trained behavioral trials in the corridor assay (**Figure 5A**). Compared to non-LED control trials, inhibiting PRNc glutamatergic activity led to markedly reduced BW performance (**Figure 5B–C**), with the average velocity decreased by over 50% (**Figure 5F**) and the time to exit the corridor more than doubled (control: 3.93 ± 0.29 s; inhibition: 9.65 ± 1.10 s, *p* = 0.003, paired t-test, *n* = 5 mice). This was mainly due to the increased number of pauses during backing out of the corridor (**Figure 5G**). In addition, stride length was reduced by the inhibition (**Figure 5H**), suggesting that both the continuation of backward movement and the gait pattern were affected. By contrast, inhibition of PRNc glutamatergic neurons had little effect on forward locomotion (**Figure 5D-E**) except that stride length was slightly reduced (**Figure 5F-H**). Together, these results demonstrate that excitatory PRNc neurons are critical for both sustaining the momentum of backward movement and coordinating the fine temporal structure of each step, highlighting the essential role of these neurons in generating smooth and coordinated backward locomotion.

**Figure 5.**
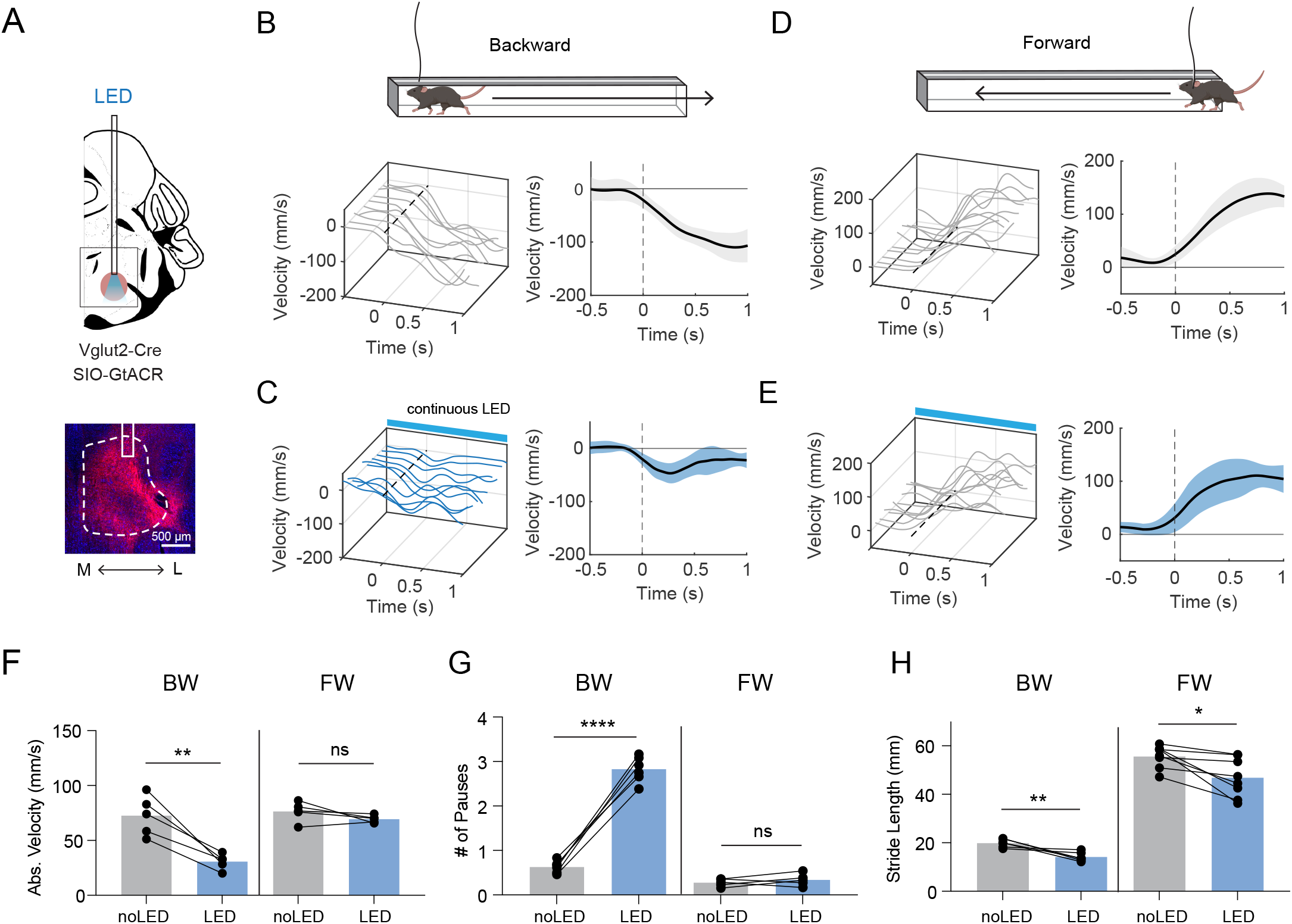
Inhibition of PRNc glutamatergic neurons impairs backward locomotion. (A) Top, schematic for viral injection and optogenetic inhibition. Scale bar = 500 µm. (B) Left, example velocity traces (gray) aligned to the onset of backward movement in the control condition. Right, mean velocity (black) ± SD (shaded) over the same 1-s window. (C) Similar to (B) but for continuous photoinhibition of PRNc vglut2 neurons. (D) Average velocity during corridor exiting. Without inhibition (LED OFF, gray) versus during continuous inhibition (LED ON, blue). **p = 0.0053, paired t-test. Lines connect individual mice (n = 5 mice). (E) Mean stride length (mm) across all limbs for LED OFF and LED ON conditions. **p = 0.0097, paired t-test. (F) Average number of pauses during corridor exiting for LED OFF and LED ON conditions. ****p < 0.0001, paired t-test.

### Anatomical connectivity of the PRNc

We next mapped the afferents of PRNc by injecting AAVretro-Cre in Ai14 reporter mice (**Figure 6A**). Retrogradely labeled (i.e. tdTomato+) neurons were observed in a wide range of structures across the forebrain and brainstem (**Figure 6B**). In the forebrain, the densest labeling was observed in the prefrontal cortex (PFC) including the anterior cingulate area (ACA), in the primary and secondary motor cortex (MOp and MOs, respectively) as well as in the primary somatosensory cortex (SSp). Neurons in these anatomically contiguous areas comprised a substantial proportion of total input cells. Significant afferents also arose from hypothalamic nuclei, particularly the zona incerta (ZI) and lateral hypothalamus (LHA). Within the midbrain, we observed robust labeling in the SC, PAG, and midbrain reticular nucleus (MRN). Nearly all of these hypothalamic and midbrain (except SC) inputs exhibited an ipsilateral bias relative to the injection site (**Figure 6C**). Additional inputs were seen from adjacent reticular regions and vestibular nuclei including the rostral pontine reticular nucleus (PRNr), GRN, IRN, medial vestibular nucleus (MV), and lateral vestibular nucleus (LAV). Finally, afferents were seen from deep cerebellar nuclei, including the fastigial nucleus (FN) and dentate nucleus (DN). Overall, PRNc receives broad inputs from both ipsilateral and contralateral hemispheres, with stronger ipsilateral than contralateral influences from many nuclei (**Figure 6D**).

**Figure 6.**
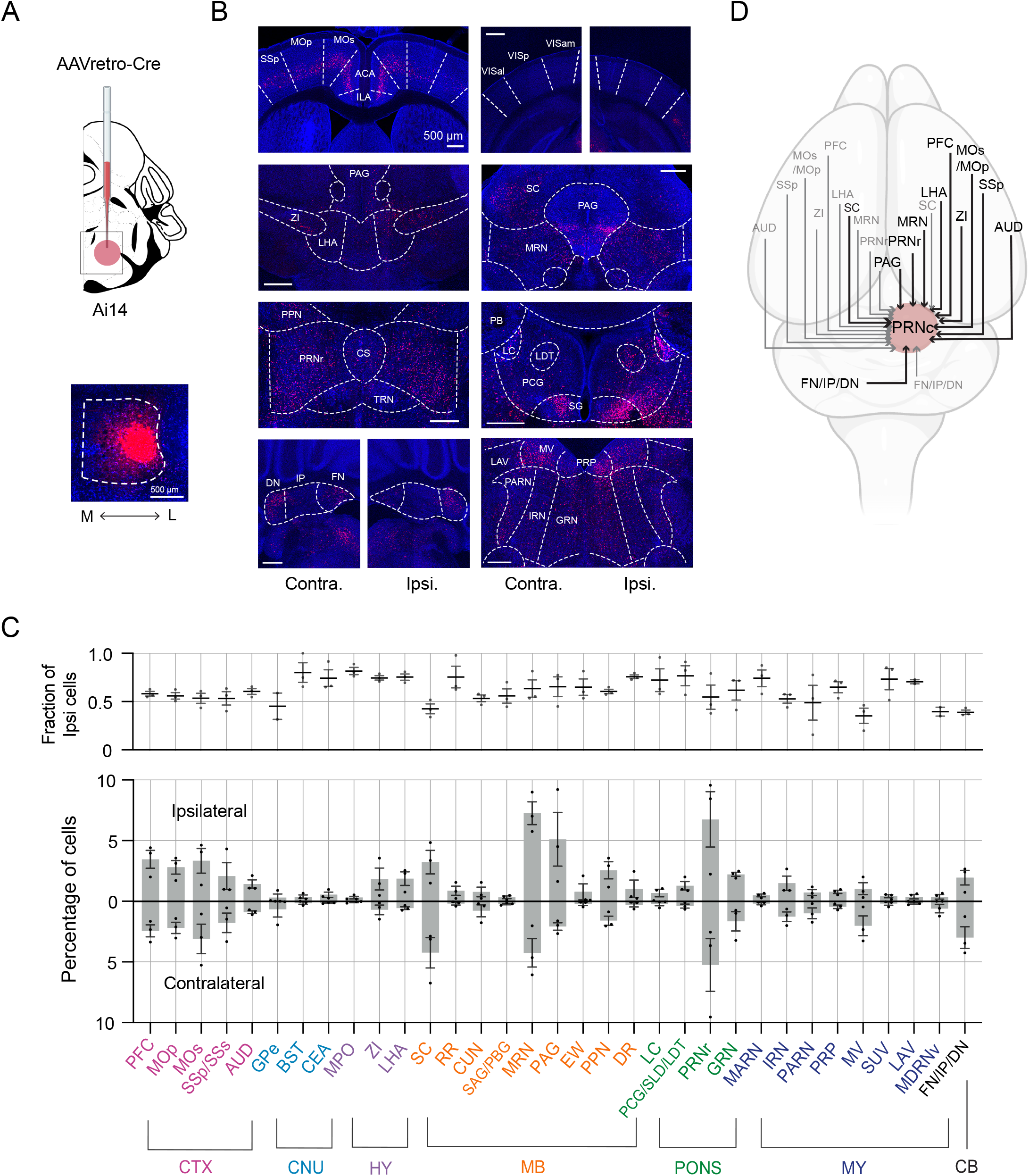
Input sources of PRNc. (A) Viral strategy and injection site for retrograde tracing. Bottom, example image of a PRNc injection site. Scale bar = 500 µm. (B) Representative images showing tdT neurons in upstream regions. From left to right and top to bottom: prelimbic (PL), infralimbic (ILA) cortices, anterior cingulate area (ACA), secondary (MOs) and primary motor cortex (MOp), primary somatosensory cortex (SSp); supplemental somatosensory area (SSs), ventral auditory area (AUDv), temporal association area (TEa); visual areas (VISam, VISp, VISal); thalamic ventromedial nucleus (VM), zona incerta (ZI), and lateral hypothalamic area (LHA); superior colliculus (SC) and periaqueductal gray (PAG); midbrain reticular nucleus (MRN); nucleus sagulum (SAG), and parabigeminal nucleus (PB); rostral pontine reticular nucleus (PRNr), superior central nucleus raphe (CS), and trigeminal motor nucleus (TRN); pontine central gray (PCG); nucleus prepositus (PRP), gigantocellular (GRN), intermediate (IRN), and parvicellular (PARN) reticular nucleus, medial vestibular nucleus (MV), lateral vestibular nucleus (LAV), and superior vestibular nucleus (SUV); fastigial (FN), interposed (IP), and dentate (DN) cerebellar nuclei. Dashed outlines mark each structure. All scale bars = 500 µm. (C) Quantification of input distribution. Bars show mean ± SEM of the percentage of total tdT cells in each region, plotted for ipsilateral (top) and contralateral (bottom) hemispheres (n = 3 mice). Regions are color-coded by major division: cortex (CTX), cerebral nuclei (CNU), hypothalamus (HY), midbrain (MB), pons, medulla (MY), and cerebellum (CB). (D) Schematic summary of PRNc afferents. Arrow thickness reflects the relative strength of inputs from each region, with the thickest lines indicating the most prominent sources.

Moreover, we characterized cell-type specific efferent projections from PRNc glutamatergic neurons by injecting Cre-dependent anterograde tracers in Vglut2-Cre mice (**Figure 7A**). Major terminal fields appeared in thalamic nuclei: the mediodorsal nucleus (MD) and parafascicular nucleus (PF) (**Figure 7B**). Within the midbrain, dense projections returned to the ZI, SC, MRN and PAG, forming reciprocal loops with these structures. These efferents to thalamic and midbrain targets usually exhibited a contralateral bias (**Figure 7C**). Additionally, robust efferent labeling extended into more caudal brainstem reticular nuclei, the GRN and medial dorsal reticular nucleus (MDRN). These projections are predominantly ipsilateral and likely continue to comprise a major descending pathway toward the spinal cord. In line with this projection, axonal labeling was also observed in the cervical spinal cord (**Figure 7B**), predominantly on the ipsilateral side (**Figure 7D**).

**Figure 7.**
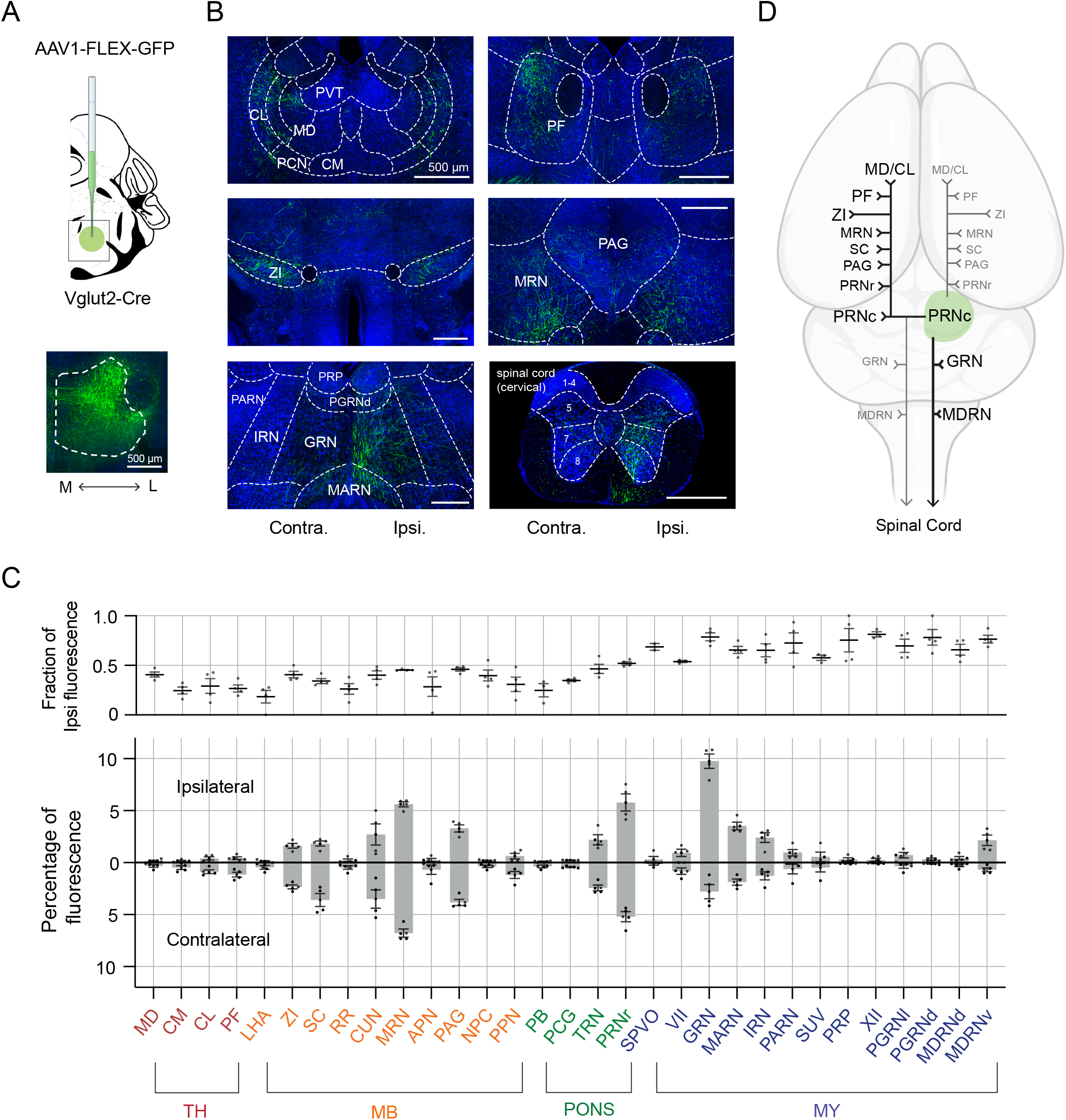
Output regions of PRNc glutamatergic neurons (A) Viral strategy and injection site for anterograde tracing. Bottom: image of a PRNc injection site (GFP, green; Nissl, blue). Scale bar = 500 µm. (B) Representative images showing GFP axon terminals in multiple downstream targets. From left to right and top to bottom: mediodorsal thalamic nucleus (MD); parafascicular thalamic nucleus (PF); zona incerta (ZI); periaqueductal gray (PAG); midbrain reticular nucleus (MRN), red nucleus (RN), rostral pontine reticular nucleus (PRNr) and retrorubral area of the midbrain reticular nucleus (RR); trigeminal motor nucleus (TRN); hypoglossal nucleus (XII), gigantocellular (GRN), intermediate (IRN), parvicellular (PARN), and medullary (MDRN) reticular nucleus. Dashed outlines mark each structure. All scale bars = 500 µm. (C) Quantification of projection strength. Bars show mean ± SEM of the percentage of total GFP labeling in each region, plotted for ipsilateral (top) and contralateral (bottom) hemispheres (n = 3 mice). Regions are color-coded by division: thalamus (red), midbrain (orange), pons (green), and medulla (dark blue). (D) Schematic summary of PRNc efferents. Arrow thickness reflects relative projection strength.

## Discussion

In this study, we identified the caudal pontine reticular nucleus (PRNc) as a previously unrecognized neural center specifically controlling backward locomotion. Combining a newly established behavioral paradigm with various functional and anatomical approaches, we show that PRNc glutamatergic neurons play an important role in selectively encoding backward movement and are sufficient and necessary for initiating and sustaining backward walking.

These findings address a key question for our understanding of direction-specific locomotor control and provide a framework to systematically investigate the neural substrates underlying locomotor coordination.

### Backward locomotion: functionally important but underexplored

Although forward locomotion has been extensively studied ^2–7^, the neural basis of backward movement has remained elusive largely due to its context-dependent nature and the difficulty of eliciting it under standard laboratory conditions. In natural environments, animals engage in backward locomotion primarily in highly specific scenarios, such as navigating through constrained spaces where turning is not feasible or retreating from danger when maintaining visual contact with the source is critical. Our behavioral observations are consistent with this general understanding: mice rarely initiate backward walking in open arenas, reinforcing the idea that this mode of locomotion is selectively deployed and not part of the animal’s default motor repertoire.

This behavioral specificity may partly explain why backward locomotion has received less attention in systems neuroscience compared to forward locomotion. Unlike forward walking which can be readily observed and measured across a range of spontaneous and task-driven behaviors, backward movement typically requires specific environmental constraints or stimuli to be reliably triggered. As a result, incorporating backward locomotion into standard locomotion paradigms has been challenging. Moreover, because backward movement is often brief, transitional, or embedded within defensive responses, it has often been considered a secondary aspect of motor behavior rather than a behavior governed by dedicated neural substrates.

Yet, from an ethological perspective, backward movement appears evolutionarily conserved and may contribute importantly to flexible, goal-directed action. It allows animals to maintain sensory contact with a threat while increasing physical distance, to navigate through spatially constrained environments, and to strategically reposition to attack a prey. The fact that backward locomotion is preserved across species ranging from insects and fish to mammals, suggests the existence of specialized neural control mechanisms, even if they are engaged only under highly specific behavioral states or environmental conditions. By establishing a reliable assay for inducing backward movement through training, we were able to overcome these limitations and systematically investigate its neural underpinnings.

### Differential control of backward and forward movement

An increasing body of evidence across species supports the notion that forward and backward locomotion are controlled by partially segregated mechanisms rather than a single, bidirectional motor program. One major line of evidence stems from comparisons of muscle activation patterns. Although some muscle synergies are shared, electromyographic (EMG) recordings reveal marked differences in both timing of activation peaks and intensity of muscle activity between forward and backward locomotion ^26,35–37^. These differences in muscle output are paralleled by distinct movement kinematics. Our kinematic analyses similarly demonstrated that backward locomotion is associated with slower average speed, smaller stride length, and higher step frequency compared to forward locomotion, consistent with previously reported findings^9,25–27,38^. Additionally, detailed kinematics studies have found that joint angle trajectories, particularly at the knee and ankle, are significantly different ^25^. These findings collectively suggest that backward locomotion is not a simple reversal of forward locomotion.

At the spinal cord level, forward and backward locomotion are associated with overlapping yet distinct patterns of neuronal recruitment. For instance, a recent study in decerebrate cats identified interneuron subpopulations that are active during both forward and backward stepping, as well as those that are selectively recruited in a direction-specific manner ^12,8^. Additionally, backward locomotion can be selectively triggered by epidural stimulation targeted to spinal segments L6–L7 in cats ^39^. Together, these findings support the existence of both direction-independent and direction-specific spinal locomotor networks.

This concept is further supported by findings in other species. For instance, distinct neuronal populations selectively activated during backward movement have been identified in fruit flies ^10,11^ and lampreys ^12^. Despite these findings, it has remained unclear whether mammals possess dedicated circuits for backward movement. Our results here suggest that PRNc may represent a specialized brainstem command module within the reticular formation involved in the control of backward locomotion.

### Functional roles of PRNc

The pontine reticular nucleus (PRN) is a component of the broader reticular formation, a diffuse network of interconnected neurons that extends from the midbrain down to the spinal cord ^40^.

The PRN is a centrally located structure in the pons and is divided into rostral (PRNr) and caudal (PRNc) portions based on its broad rostrocaudal span. Despite its prominent anatomical position, the functional role of the PRNc has remained underexplored compared to neighboring nuclei.

PRNc has previously been suggested in mediating acoustic startle responses (ASRs) ^36–38^. Specifically, “giant neurons” in the ventrolateral PRNc receive direct input from cochlear root neurons and project monosynaptically to spinal motor neurons, with lesions or direct stimulation of this region abolishing ASRs or eliciting ASR-like responses, respectively ^41–44^. Additionally, the PRNc contributes to multisensory processing, e.g. transmitting auditory and other sensory information to the PCG (a relatively minor projection; **Figure 7**) and ultimately modulating higher-order limbic regions through a reticulo-limbic pathway ^45,46^.

Our results significantly extend the known functional repertoire of PRNc by demonstrating its selective involvement in backward locomotion. Thus, PRNc neurons appear capable of mediating at least three distinct functional roles: rapid reflexive responses via direct spinal cord projections, emotion-related functions via multisensory projections to higher-order limbic structures, and voluntary motor behaviors through broader network integration. This functional diversity raises the possibility that different subsets of PRNc neurons collectively contribute to coordinating defensive reflexes, sensory integration, and direction-specific locomotion. In the future, it would be of interest to determine whether these different functional roles are mediated by overlapping or distinct PRNc subgroups.

### Input-output organization of PRNc

#### Inputs of PRNc

Our anatomical tracing positions the PRNc as a central integrator of diverse signals that may coordinate backward locomotion. Inputs to the PRNc can be broadly categorized into cortical, defense-related, and locomotor-related sources. Cortical inputs include projections from areas associated with motor planning and sensory integration, such as the prefrontal cortex, primary and secondary motor areas, and somatosensory cortex. These inputs may support the heightened attentional demands required to process sensory input, evaluate contextual information, and refine motor control during backward locomotion, consistent with evidence of increased cortical engagement during backward walking ^37,47^. The PRNc also receives robust input from hypothalamic and midbrain regions implicated in defensive behaviors, including LHA, ZI, PAG and SC ^31,32,48–50^. These connections align with employing backward movement in defensive contexts, highlighting a potential role of PRNc in processing arousal-and threat-related signals. In addition, PRNc receives afferents from established locomotor centers, including the pedunculopontine nucleus (PPN), cuneiform nucleus (CUN), and GRN ^13,14,16,51–53^, suggesting its integration within a broader motor command network. While the functional relationship between PRNc and these locomotor regions remains unclear, their convergence onto PRNc may help integrate contextual, sensory, and motor inputs to coordinate precisely timed and context-appropriate backward movements.

#### Projections from PRNc

Beyond its role as an integrative input hub, glutamatergic neurons in the PRNc project to several key regions. They send ascending projections to the MD and PF thalamic nuclei, which are known for relaying motor planning signals to the prefrontal cortex and modulating basal ganglia circuits, respectively ^54–59^. This suggests a pathway by which PRNc could modulate higher order motor or sensory systems during backward locomotion, supporting the idea of parallel processing of movement and sensory state ^13^. Dense projections are also seen in the PAG, ZI and SC, which notably are major sources of input to PRNc. This reciprocal connectivity raises the possibility of a feedback loop that could help with continuous integration of sensory and threat-related information, potentially influencing the timing and execution of backward movement.

The most prominent descending projections of PRNc pass through reticular nuclei, including GRN and MDRN, ultimately targeting the spinal cord, consistent with projection patterns of reticulospinal neurons ^60–63^. Similar descending pathways from other locomotor populations in the brainstem, such as those controlling forward running, stopping, turning, and forelimb movement, also directly or indirectly recruit spinal motor networks as the final effectors of movement ^15,17,19,52,64^. Further clarification of how PRNc projections interact with spinal networks to influence backward locomotion may provide valuable insight into the broader organization of direction-specific motor circuits.

Taken together, our findings identify glutamatergic neurons in the PRNc as a critical neuronal population governing backward locomotion. These neurons may integrate diverse arousal, defensive, and motor-control inputs to coordinate motor outputs. This study may also advance our understanding of the modular organization of the brainstem and direction-specific locomotor control. Future investigations of additional neuronal populations within the PRNc, such as inhibitory neurons and their interactions with glutamatergic populations, will further clarify the neural mechanisms underlying backward locomotion. Furthermore, these findings may provide insights relevant to exploring therapeutic strategies for neurological disorders characterized by directional movement deficits, gait disturbances, or impaired motor coordination, such as Parkinson’s disease, stroke, and spinal cord injuries.

## Acknowledgements

This work was supported by grants from the US National Institutes of Health (DC008983, DC020887, NS132912 to L.I.Z.; EY019049, MH116990 and AG089756 to H.W.T.).

## Author Contributions

M.B.S. performed most of the behavioral, imaging and anatomical experiments and data analysis.

J.H. and G.W.Z. helped with the calcium imaging experiments and data analysis. H.W.T and L.I.Z. supervised the project. M.B.S., L.I.Z., and H.W.T. wrote the manuscript.

## METHODS

All experimental procedures in this study were in accordance with the guidelines for the care and use of laboratory animals of the US National Institutes of Health (NIH), and were approved by the Animal Care and Use Committee (IACUC) of the University of Southern California.

### Animals

Mice used in this study included C57BL/6J, Trap2 (JAX strain # 030323), and Ai14 (Cre-dependent tdTomato reporter, JAX strain # 007914), Vglut2-Cre (JAX strain # 016963), obtained from Jackson Laboratory. Both male and female adult mice (2–6 months old) were used and housed with a 12 h light-dark cycle, at 65–75°F temperature and 40–60% humidity.

### Viral constructs and purpose

AAV1-EF1a-FLEX-GCaMP7s [1.6 × 10^13^GC/ml, Addgene] was used for cell type-specific fiber photometry recording in Vglut2-Cre mice. AAV1-EF1a-DIO-hChR2-eYFP [1.82 × 10^13^ GC/ml, Addgene] was used for cell type-specific activation in Vglut2-Cre mice. pAAV-hSyn1-SIO-stGtACR2-FusionRed [1.9 × 10^13^ GC/ml, Addgene] was used for cell type-specific inhibition in Vglut2-Cre mice. AAV1-CAG-FLEX-GFP-WPRE [2 × 10^13^ GC/ml, Addgene] was used for cell type-specific anterograde tracing in Vglut2-Cre mice. Due to the low labeling efficiency observed with cell-type-specific pseudorabies retrograde tracing from PRN regions in our experiments (data not shown), which was consistent with previous reports ^65–67^, we instead used AAVretro-Cre [1.5 × 10¹ GC/ml, Vigene] to broadly map inputs to PRNc neurons.

### Viral injection

Mice were anesthetized with 1.5–3% isoflurane inhalant and placed on a stereotaxic apparatus with a heating pad (37.5 °C) in a sterile surgical hood. Buprenorphine and ketoprofen were subcutaneously injected as analgesics. To expose the brain surface, a single incision was made on the skin above the skull. Using stereotaxic coordinates (coordinates: AP –5.26 mm, ML 0.8 mm, DV –5.00 mm), the injection site was marked unilaterally on the skull. A small burr hole (<300 μm) was drilled, and any remaining skull was removed to expose the dura. The site was then flushed with sterile saline. A beveled glass pipette was loaded with 2 μL of AAV encoding various proteins (details listed above) depending on the experimental purpose and mouse strain.

The pipette was positioned over the hole and lowered through the dura to the target depth using a micromanipulator. A volume of 60 nL was injected at a slow rate (15–25 nL/min). After injection, the pipette was left in place for 5–10 minutes before being retracted. The skin was sutured with sterile 4-0 nylon thread. Mice recovered for at least one week before further procedures.

### Fiber implantation

For fiber photometry or photostimulation experiments, 2–3 weeks after the viral injection, an optic cannula (200 µm core, NA= 0.22, RWD Inc.) was implanted 300 µm above the injection site. The procedure followed the same steps as the viral injection until the pipette insertion stage. Instead of a pipette, the optic cannula (200 μm diameter) was slowly inserted using a micromanipulator (coordinates: AP –5.26 mm, ML 0.8 mm, DV –4.70 mm). Once positioned at the intended depth, dental cement was applied around the cannula for mechanical stability. The mice were allowed to recover for at least 1 week before the behavior tests. After experiments, the brain was extracted, sectioned, and imaged to confirm the locations of implanted cannulas.

### Training of backward walking in corridor

Prior to the experiments with fiber photometry and optogenetic inhibition, mice underwent backward-walking training following a 16–18 h water deprivation. Training took place in an open-field arena housing a clear acrylic corridor (4 cm W × 30 cm L × 4 cm H) with a 1 cm reward port on one of the distal ends. At the start of each trial, the mouse entered the corridor and, upon reaching the closed end, received a 30 uL water reward through the port. Once the mouse entered and reached the end of the corridor, a water droplet was given through the opening. Exiting the corridor completely was required to trigger the next trial and obtain another reward. Training continued for 5–7 days or until mice reliably executed ≥30 backward bouts per session (about 20-30 minutes per session).

For the c-Fos trapping experiment, a separate acrylic corridor was designed to have both ends opened, and also to enable water reward at the middle of the corridor. The dimension of the corridor remains the same, but with an additional 1-cm opening at the top middle made. Animals were water deprived the day before the training began. For the first 15 minutes, the corridor was placed at the center at one wall of the arena so that the animal could exit at the opposite end of where they entered. Once they entered, the water droplet was given at the center top opening and the animal would exit through the other open end. For the next 15 minutes, the corridor was pushed to the corner so that one end is now closed. Once they entered, the animals would have to reach the closed end to get the water droplet reward and then back out of the corridor to exit in order to initiate a new trial.

### Behavioral annotations

Behavioral videos were from top-down, bottom-up, or both depending on the experiment. Video was recorded at 60 frames per second (fps) using a GoPro under consistent lighting (GoPro HERO10 Black). Pose estimation was conducted with DeepLabCut (DLC) v3.0.0, a deep learning framework for markerless tracking. Key body parts (nose, neck, tail base, and all limbs) were manually labeled in extracted frames to train a ResNet-50-based model. The trained network generated x-y coordinates and likelihood values. Tracking data were interpolated for low-likelihood detections (threshold: 0.6). Processed coordinates were cleaned in MATLAB and used to quantify behavioral metrics, including velocity, angular velocity, head angle, and gait patterns.

### Fiber photometry calcium imaging

Cre-dependent GCaMP7s virus was injected into the PRNc in Vglut2-Cre mice followed by an optic fiber implantation. After 2 weeks, the calcium signal was recorded using the RWD Fiber Photometry System (R821 Tricolor Multichannel). Following 5–7 days of behavioral training, animals were put into the same behavior arena with a narrow slit in the ceiling to allow for fiber movement. The implanted fiber was connected to the RWD Fiber Photometry System (R821 Tricolor Multichannel). Excitation was delivered via a 470 nm LED (GCaMP7s signal) and 410 nm LED (reference) and sampled at 60 fps through the OFRS software. The animals received a random trial water reward of 30 μL after reaching the end of the corridor. Water rewarded trials (30 µL at corridor end) were excluded from analysis to avoid reward related transients. Data were obtained and analyzed using custom MATLAB software.

### Ca2+ dynamics analysis

Calcium signals were analyzed using customized MATLAB scripts. To correct for motion artifacts and non-calcium-related fluctuations, we used the 405 nm signal as a reference for the calcium-dependent 490 nm signal. A linear regression was performed to fit the 405 nm signal to the 490 nm trace, and this fitted signal was used to calculate ΔF/F using the following equation:

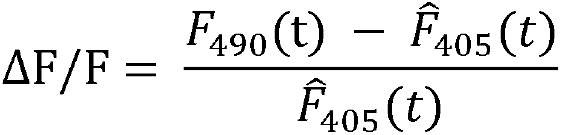

Where Fᶺ_405_ (*t*) is the fitted 405 nm signal. We truncated a 3-s window around movement onset (1 s before to 2 s after) and converted to z-scores using

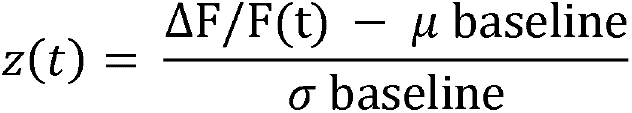

where the baseline refers to the 1 s pre-onset period, and μ and σ are the mean and standard deviation of ΔF/F during that window.

### Calcium-velocity correlation

To assess the relationship between PRNc calcium activity and locomotor speed, we applied a sliding window analysis. From the z scored ΔF/F and velocity traces spanning 0–2 s after movement onset, we extracted overlapping windows (10 frames, stepped every 5 frames). Within each window, we calculated the mean calcium response and mean velocity. We then pooled these windowed means across all forward and backward trials and computed the Pearson’s correlation coefficient (*r*) and its p-value.

### c Fos TRAP2 labeling of PRNc activation

To permanently label neurons activated during backward versus forward locomotion, we used TRAP2::Ai14 transgenic mice. On the test day, mice received an intraperitoneal injection of 4 hydroxytamoxifen (4 OHT, 50 mg/kg) dissolved in corn oil. Immediately following the injection, animals were placed into the corridor apparatus under one of two conditions: 1) Forward only (FW) group: mice entered through one of two open-ends and received a 30 µL water reward at that central port and exited through the other open end. 2) Backward (BW) group: mice entered through the open end and received a 30 µL reward at the closed end port and then backed out to exit through the same entrance. Each session continued until mice completed 50 rewarded trials (forward in FW, backward in BW). Individual animals were then single housed, and were deeply anesthetized and perfused after 10 days for brain harvesting to visualize TRAPed tdT+ neurons.

### RNAscope

A TRAP2::Ai14 double transgenic mouse line was used to label neurons activated during behavior. Ten days following the c-Fos trapping procedure, animals were deeply anesthetized and transcardially perfused with 4% paraformaldehyde (PFA) in phosphate-buffered saline (PBS). Brains were post-fixed overnight in 4% PFA at 4°C, then dehydrated in a series of sucrose solutions (10%, 20%, and 30%). Coronal sections (25 μm thick) were cut using a cryostat and mounted on Superfrost Plus microscope slides (Thermo Fisher Scientific).

RNAscope staining was performed using the RNAscope® Multiplex Fluorescent Reagent Kit v2 (Advanced Cell Diagnostics, #323100), according to the manufacturer’s protocol. Probes targeting *tdTomato* (to label TRAPed c-Fos+ neurons) and *Slc17a6* (Vglut2) were used for co-localization analysis. All probes were designed and validated by Advanced Cell Diagnostics.

Following hybridization and amplification, sections were cover slipped and imaged using confocal microscopy.

### Optogenetic manipulation

For all experiments, optic fibers implanted in the animals were connected to patch cables via ceramic sleeves. Mice were placed in a custom behavioral corridor with a transparent acrylic floor (4 cm W × 50 cm L × 4 cm H), allowing video acquisition from below at 60 fps. In animals expressing channelrhodopsin-2 (ChR2), photostimulation was delivered using a 470 nm LED (Thorlabs) controlled via a custom LabVIEW interface. Light pulses (5 mW) were administered for 1 second at either 5 Hz, 10 Hz, or 20 Hz. Stimulation occurred only when the animal was positioned such that there was sufficient space behind it in the corridor to allow for potential backward movement. A minimum of 20 stimulation trials were collected per animal. For GtACR-expressing animals, photoinhibition was achieved by continuous 470 nm LED illumination delivered as the animal entered the corridor from an adjacent open field. The light remained on for the duration of the animal’s presence in the corridor and was delivered in a pseudorandomized fashion, allowing for interleaved control (LED-off) and stimulation (LED-on) trials. At least 15 LED-on and 15 LED-off trials were collected per animal.

### Histology

Mice were anesthetized with isoflurane and perfused with 4% PFA and 1x PBS. Brains were dissected out and fixed overnight in 4% PFA at 4°C. After fixation, tissues were embedded in agarose and sectioned at 150 µm using a vibratome (Leica, vt1200s). Sections were nissl stained overnight and mounted and cover slipped on superfrost glass slides.

### Image acquisition

Fluorescent images were acquired using an Olympus FV1000 confocal laser scanning microscope with 4×, 10×, and 40× objectives, selected based on the required resolution and anatomical detail. Channels included eYFP/GFP (excitation: 488 nm; emission: 500–550 nm), tdTomato (excitation: 543 or 561 nm; emission: 570–620 nm), and the 405 Nissl stain (excitation: 405 nm; emission: 420–480 nm). All imaging parameters, including laser power, gain, and detector offset, were optimized to avoid signal saturation and held constant across samples within each experimental group to ensure comparability. Images were acquired as single-plane optical sections and exported using Olympus FV Viewer or Fiji/ImageJ for subsequent analysis and figure preparation.

### Quantification and Statistical analysis

#### QUINT Pipeline for Cell Counting

Quantification of labeled neurons across brainstem regions was performed using the QUINT pipeline ^68^, a semi-automated workflow for atlas registration and cell counting. First, histological images were aligned to the Allen Mouse Brain Atlas using DeepSlice ^69^, an AI-based tool for identifying the best-matching atlas sections. The output was then refined using VisuAlign to manually adjust atlas overlays and ensure accurate registration of brainstem subregions. Following registration, fluorescently labeled cells were segmented using Ilastik, a machine learning-based image segmentation tool. A pixel classification workflow was trained to differentiate labeled cells from background signals. The resulting probability maps were exported and further processed with Nutil, which was used for region-based cell counting.

#### Projection intensity quantification

To quantify projection intensity in specific brainstem targets, images were first registered to the Allen Brain Atlas using the same DeepSlice and VisuAlign workflow described above. Following atlas alignment, the oib image files were imported into Fiji/ImageJ. Using the ROI Manager, ROIs were manually drawn over target regions based on Visualign registration of subregions. For each ROI, a threshold was applied to the channel of interest to eliminate background fluorescence, ensuring that only the true signal was measured. Within each ROI, the area of above-threshold pixels (i.e., signal-positive area) was quantified using the “Measure” function in Fiji. To enable comparison across animals and groups, the signal-positive area in each ROI was normalized to the total signal-positive area across all ROIs for that section, yielding a percentage of total projection signal per region.

#### Pose estimation

Pose estimation was performed using DeepLabCut ^20,21^ (version 3.0), a deep learning-based tool for markerless tracking of user-defined body parts. A custom-labeled training dataset was created to annotate key anatomical landmarks, including the nose, neck, tail base, and the center of each of the four paws. The network was trained using the ResNet-50 architecture on the PyTorch framework until pose estimation accuracy plateaued, with no further performance improvements observed. Following the training, the predicted body part coordinates were used for analysis using custom MATLAB scripts for following behavioral and kinematic analyses.

#### Quantification of velocity

For velocity analysis, the tail base was used as the primary tracking point to estimate locomotor speed. The neck-to-tail base vector defined the body’s longitudinal axis and served as the reference for determining movement direction for each frame.

Instantaneous velocity was calculated as the frame-to-frame displacement of the tail base. To assign directionality, the velocity vector was projected onto the body axis: positive values indicated forward movement (toward the neck), while negative values indicated backward movement (away from the neck). These directional velocity values were used for further behavioral segmentation and kinematic analysis.

#### Gait analysis

For gait analysis, stance phases for each limb were annotated manually using a custom MATLAB interface. Stance onset was defined based on visual confirmation of full paw contact with the floor. Based on these annotations, key gait parameters were calculated, including stride length—defined as the distance between consecutive stance onsets of the same paw—and step frequency, measured as the number of steps per unit time for each limb.

## Statistics

All statistical analyses were performed using GraphPad Prism (version 8). Significance levels are reported as follows: *\*P < 0.05, **P < 0.01, ***P < 0.001, ****P < 0.0001*. Shaded areas in line plots represent the mean ± standard deviation (SD), unless otherwise specified.

Error bars denote the mean ± standard error of the mean (SEM), unless otherwise noted. Sample sizes were determined based on prior pilot experiments or from related research (reference).

Animals were randomly assigned to experimental and control groups to minimize potential bias. Normality of data distributions was assessed using the Shapiro–Wilk test. For comparisons involving more than two groups, one-way or two-way analysis of variance (ANOVA) was performed, followed by Tukey’s or Bonferroni post hoc tests for multiple comparisons, as appropriate. For comparison of variability, pairwise F-test was used. For comparisons between two independent groups with normally distributed data, an unpaired two-tailed *t*-test was used. For within-subject comparisons (e.g., pre-vs post-manipulation from the same animal), a paired *t*-test was applied. Exact *P*-values, test statistics, and degrees of freedom are reported in the relevant figure legends.

